# GraFusionNet: Integrating Node, Edge, and Semantic Features for Enhanced Graph Representations

**DOI:** 10.1101/2024.11.22.624875

**Authors:** Md Toki Tahmid, Tanjeem Azwad Zaman, Mohammad Saifur Rahman

## Abstract

Understanding complex graph-structured data is a cornerstone of modern research in fields like cheminformatics and bioinformatics, where molecules and biological systems are naturally represented as graphs. However, traditional graph neural networks (GNNs) often fall short by focusing mainly on node features while overlooking the rich information encoded in edges. To bridge this gap, we present **GraFusionNet**, a framework designed to integrate node, edge, and molecular-level semantic features for enhanced graph classification. By employing a dual-graph autoencoder, GraFusionNet transforms edges into nodes via a line graph conversion, enabling it to capture intricate relationships within the graph structure.

Additionally, the incorporation of Chem-BERT embeddings introduces semantic molecular insights, creating a comprehensive feature representation that combines structural and contextual information. Our experiments on benchmark datasets, such as Tox21 and HIV, highlight GraFusionNet’s superior performance in tasks like toxicity prediction, significantly surpassing traditional models. By providing a holistic approach to graph data analysis, GraFusionNet sets a new standard in leveraging multi-dimensional features for complex predictive tasks.

**CCS CONCEPTS:**

- **Computing methodologies → Neural networks**.

**ACM Reference Format:** Md Toki Tahmid, Tanjeem Azwad Zaman, and Mohammad Saifur Rahman. 2018. GraFusionNet: Integrating Node, Edge, and Semantic Features for Enhanced Graph Representations. In *Proceedings of Make sure to enter the correct conference title from your rights confirmation email (Conference acronym ‘XX)*. ACM, New York, NY, USA, 9 pages. https://doi.org/XXXXXXX.XXXXXXX

## 1 INTRODUCTION

The advent of graph neural networks (GNNs) has ushered in a new era in the realm of data science, particularly in fields where data can be naturally represented as graphs. This is especially true in cheminformatics and bioinformatics, where molecules and biological structures are inherently graph-like, with atoms as nodes and bonds as edges. Traditional machine learning approaches often struggle to capture the rich, relational information embedded within these structures, limiting their ability to predict properties and behaviors of complex molecules accurately. Our research introduces a novel dual-graph autoencoder framework that not only captures the intricate relationships between nodes and edges but also provides a generalized approach applicable to any dataset representable in a graph format. This dual approach is designed to leverage both significant node and edge features, offering a more nuanced and comprehensive understanding of the underlying graph structure.

Recent literature on graph neural networks has explored various aspects of their application in chemical and biological data analysis. Kipf and Welling’s work on semi-supervised classification with graph convolutional networks (GCNs) has laid the groundwork for subsequent developments in the field [7]. Additionally, the concept of graph autoencoders has been further explored by Simonovsky and Komodakis, who introduced a novel neural network architecture for graph-based data [13]. Despite these advances, the exploration of dual-graph representations, which encode both node and edge information to enrich the feature space, remains relatively untapped. Our approach seeks to fill this gap by integrating a dual-graph autoencoder with Chem-Bert embeddings, demonstrating its applicability across a wide range of graph-based data analysis tasks, extending the insights provided by Hamilton et al. in their comprehensive review on graph representation learning [4].

Our contribution to the advancement of graph-based analytics is significant. The core of our innovation lies in the dual-graph autoencoder architecture, which ingeniously combines the latent representations of both the original and dual graphs, capturing a holistic view of the molecular or structural data. This method not only enriches the feature set by incorporating both atom and bond characteristics but also enhances model interpretability and predictive performance. Our empirical evaluations across diverse datasets underscore the versatility and effectiveness of our approach, marking a step forward in the application of GNNs to complex datasets. By demonstrating the general applicability and superior performance of our dual-graph autoencoder framework, we pave the way for future explorations and applications in various domains where graph-based data representation is paramount.

**Figure 1:**
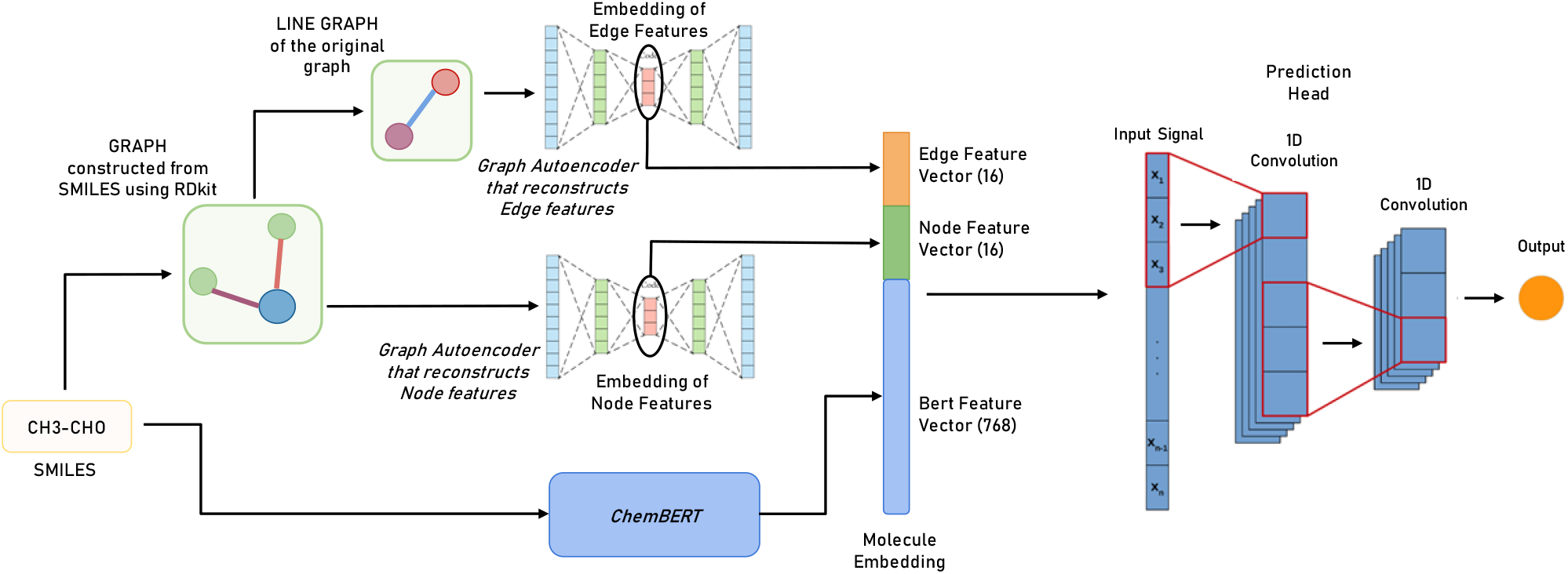
Complete Architecture of the Dual-Graph Autoencoder Model.

## 2 RELATED WORK

Graph Neural Networks (GNNs) have emerged as a transformative approach to solving complex molecular tasks, particularly in datasets like Tox21 and HIV. These datasets pose significant challenges due to the intricate relationships between molecular structures and their properties. Studies such as [6] and [9] demonstrated the ability of GNNs to model these complex interactions effectively, outperforming traditional descriptor-based methods. These works underscored the potential of GNNs to predict molecular toxicity, drug interactions, and other critical chemical properties with high accuracy. Additionally, advancements by [14] highlighted the importance of incorporating edge features, such as bond types and distances, into GNN architectures. These features significantly improved the predictive performance by better capturing the nuances of molecular graphs. Together, these studies demonstrate that GNNs, by leveraging graph-based representations, are revolutionizing the field of molecular modeling and analysis [3, 15]. Graph autoencoders (GAEs) have also played a pivotal role in the domain of molecular feature extraction, providing an unsupervised mechanism to encode the complex structural and chemical information inherent in molecular graphs. The work by [8] introduced GAEs as an effective tool to enhance the representation of molecular features, particularly in property prediction tasks. Their model leveraged graph embeddings to capture detailed relationships between atoms and bonds, facilitating improved downstream analysis. The incorporation of variational techniques in GAEs, as explored by [10], expanded their application by generating chemically valid molecular graphs while preserving critical properties. These advancements highlight how GAEs can transform raw molecular data into meaningful representations, enabling more accurate predictions in tasks like drug discovery and toxicity analysis [1, 11].

The integration of edge features in GNNs has emerged as a critical factor in improving the predictive power and interpretability of these models. By explicitly encoding bond-specific information and other edge-level attributes, researchers have been able to significantly enhance the performance of molecular prediction models. For example, the work by [2] introduced edge-based attention mechanisms, which enabled the model to weigh bond-specific features dynamically, improving the model’s ability to focus on critical molecular interactions. Similarly, the directed message-passing neural network (D-MPNN) proposed by [5] utilized edge-specific features to better represent the influence of chemical bonds in molecular systems. Furthermore, [12] emphasized the integration of physical pooling mechanisms, which combined node and edge attributes to generate more holistic molecular representations. These innovations collectively demonstrate that the inclusion of edge features is not just a technical enhancement but a necessity for achieving state-of-the-art results in molecular prediction tasks. This approach paves the way for more accurate, explainable, and robust models capable of addressing complex challenges in chemical and biological research.

## 3 MATERIALS AND METHODS

Dual-graph autoencoder uses the graph’s node features and edge features in a combined feature space. We first use the graph’s orginal node features to embed the graph into a fixed dimensional space. With the idea of dual graph conversion, the edge features are converted to node features. This converted graph’s feature space allows us to encode the edge properties into a fixed dimension. The concatenated feature space allows us to use both the node and the edge features during downstream tasks. In this section we describe the datasets we used, feature representation of the molecules, and the architecture of dual-graph autoencoder.

**Table 1:**
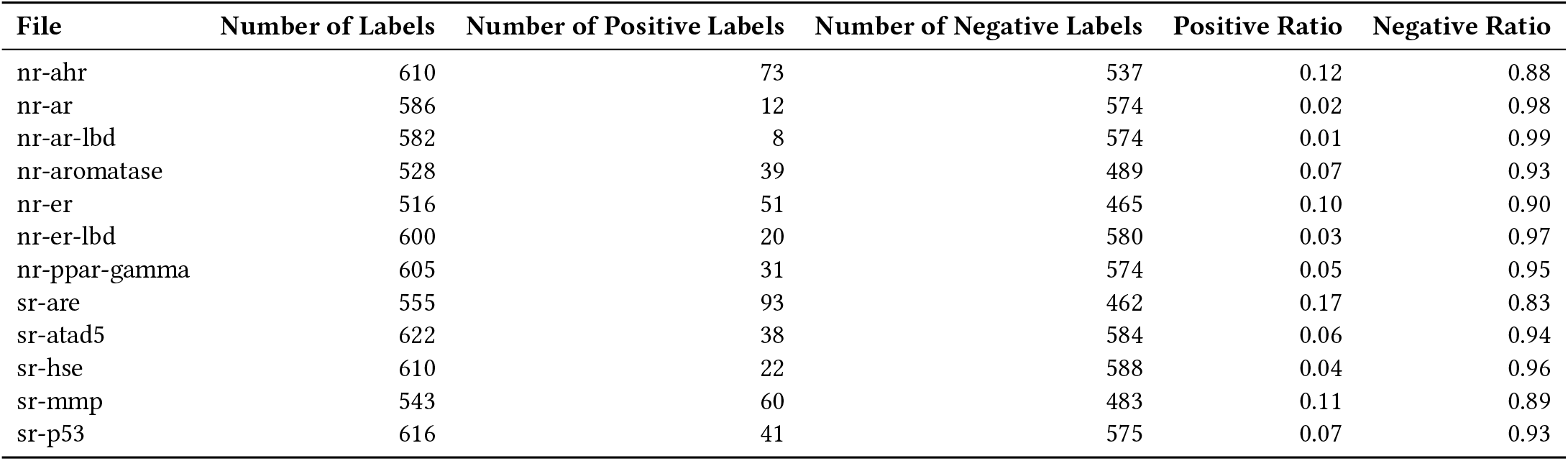
Dataset Labels Distribution.

| File | Number of Labels | Number of Positive Labels | Number of Negative Labels | Positive Ratio | Negative Ratio |
| --- | --- | --- | --- | --- | --- |
| nr-ahr | 610 | 73 | 537 | 0.12 | 0.88 |
| nr-ar | 586 | 12 | 574 | 0.02 | 0.98 |
| nr-ar-lbd | 582 | 8 | 574 | 0.01 | 0.99 |
| nr-aromatase | 528 | 39 | 489 | 0.07 | 0.93 |
| nr-er | 516 | 51 | 465 | 0.10 | 0.90 |
| nr-er-lbd | 600 | 20 | 580 | 0.03 | 0.97 |
| nr-ppar-gamma | 605 | 31 | 574 | 0.05 | 0.95 |
| sr-are | 555 | 93 | 462 | 0.17 | 0.83 |
| sr-atad5 | 622 | 38 | 584 | 0.06 | 0.94 |
| sr-hse | 610 | 22 | 588 | 0.04 | 0.96 |
| sr-mmp | 543 | 60 | 483 | 0.11 | 0.89 |
| sr-p53 | 616 | 41 | 575 | 0.07 | 0.93 |

### 3.1 Datasets

#### 3.1.1 Basic overview of Tox21 Dataset

The Tox21 dataset is a key resource for researchers in the field of toxicology and cheminformatics, aimed at providing a comprehensive set of tools for toxicity prediction. It consists of a multi-label binary classification task involving the screening of chemicals for toxicity towards environmental and human health. This dataset includes information on a broad spectrum of chemical compounds represented by Simplified Molecular Input Line Entry System (SMILES) notations, which is a text-based representation of chemical structures allowing for easy input and sharing of molecular data.

The Tox21 challenge focuses on predicting the toxicity of compounds across 12 different assays, which are essentially biological tests to determine the presence of specific toxic effects. Each assay represents a unique label, making this a multi-label classification problem where each chemical is evaluated to ascertain whether it exhibits toxic behavior in each of the 12 assays (i.e., yielding a yes/no outcome for each label).

Originally, the dataset comprises 7,831 unique compounds, with each entry including one molecule identifier (mol_id), the corresponding SMILES notation, and the outcomes for the 12 toxicity assays. However, the dataset poses several challenges for computational models, including a significant amount of missing data (NaN values) in the labels. After further investigation, individual datasets for each of the 12 labels have been found; this aided in more efficient preprocessing for the data. Furthermore, the datasets were heavily imbalanced, consisting of significantly larger fractions of negative data points 1. Techniques such as synthetic data generation were tried, but yielded no improvements to accuracy. As such, undersampling the minority class showed the most stable results.

#### 3.1.2 Basic overview of HIV Dataset

In our work, we utilized the HIV dataset, originally introduced by the Drug Therapeutics Program (DTP) AIDS Antiviral Screen. This dataset evaluates the inhibitory potential of over 40,000 compounds against HIV replication. The screening results classify compounds into three categories: confirmed inactive (CI), confirmed active (CA), and confirmed moderately active (CM). For our study, we combined the CA and CM categories into a single “active” class, resulting in a binary classification task between inactive (CI) and active (CA + CM).

### 3.2 Feature Representation

#### 3.1.1 Feature Representation with Rd-Kit

We have used the Rd-Kit library to extract molecule level features from the smile strings. Rd-Kit provides features for each atom which we incorporate as our node feature. For the current study, we include 11 node features: atom index, atomic number, is_aromatic, hybridization, number of hydrogens, formal charge, chirality, is_in_ring, degree, implicit valence, and explicit valence. Moreover, Rd-Kit provides the facility to extract edge features between two connecting atoms. As bond features, we use: bond type, bond type as double, is_conjugated, is_in_ring, bind_dir, begin atom index, end atom index, and is_aromatic. We include the nodes and the edge features into a pytorch graph object.

#### 3.2.2 Feature Representation with Chem-Bert

In addition to graph features, we use Chem-Bert[] to extract whole molecule level feature set. Chem-Bert provides us with an extensive feature set of 768 dimensions. We concatenate the graph level features with the Bert extracted features for our final model.

### 3.3 Graph Neural Network

Graph Neural Networks (GNNs) are a class of neural networks designed to perform inference on data that can be represented as graphs. GNNs capture the dependency of graphs via message passing between the nodes of graphs.

In a typical GNN, the message passing and aggregation process can be described mathematically at each layer *l* by the following equations:

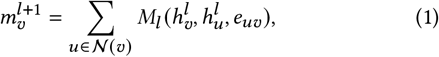

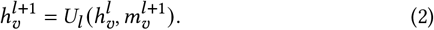

Here, 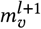 is the aggregated message for node *v* at layer *l* + 1, 𝒩 (*v*) denotes the neighboring nodes of *v, M*_*l*_ is the message function, 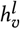 is the feature vector of node *v* at layer *l, e*_*uv*_ is the feature vector of the edge from *u* to *v*, and *U*_*l*_ is the update function.

The initial feature vector 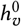 for each node is usually the node’s attributes. After *L* layers of propagation, the feature vectors 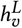 are used for downstream tasks such as node classification, link prediction, or graph classification.

### 3.4 Convolutional Graph Neural Network

Graph Convolutional Neural Networks (GCNs) are a type of neural network designed to work directly on graphs and leverage their structural information. They are particularly well-suited for node classification, link prediction, and graph classification tasks.

The core idea of GCNs is to update the representation of a node by aggregating the representations of its neighbors. This is often referred to as message passing. The following equation describes this process:

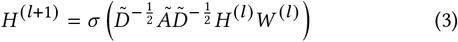

where:

- *H* ^(*l*)^ is the matrix of node features at layer *l*,
- *H* ^(*l* +1)^ is the matrix of node features at layer *l* 1,
- 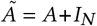 is the adjacency matrix of the graph *G* with added
- self-connections (where *I*_*N*_ is the identity matrix),
- 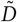is the degree matrix of 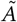,
- *W* ^(*l*)^ is the weight matrix for layer *l*,
- σ (·) is the activation function, e.g., ReLU.

The degree matrix 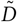 is diagonal with entries 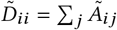. The normalization by 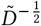 is critical to ensure that the scale of the feature representations is maintained.

GCNs generalize the convolutional operation from Euclidean data to graph-structured data. By stacking multiple such layers, GCNs can capture the hierarchical patterns in the data.

### 3.5 Graph Autoencoder

Graph autoencoders are a powerful type of neural network designed for graph-structured data. They operate by compressing the graph into a latent space (encoding) and then reconstructing it (decoding). The objective of such a model is to learn a latent representation that captures the essence of the graph’s structure.

A key part of the training process is the reconstruction loss, which often employs the Mean Squared Error (MSE) to quantify the difference between the original graph and the reconstructed graph. The MSE loss for graph autoencoders can be formulated as follows:

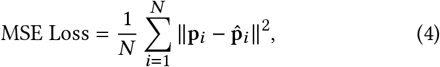

where p_*i*_ represents the original node features, and 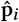 represents the reconstructed node features from the autoencoder. Minimizing this loss during training encourages the model to learn to recreate the original features as closely as possible.

### 3.6 Dual Graph Autoencoding: Incorporating both Node and Edge Features

Dual Graph Autoencoding is a novel framework designed to address the limitations of traditional graph convolutional networks (GCNs), which primarily focus on reconstructing node features while neglecting the rich information encoded in edge features. By introducing a dual encoding-decoding mechanism, this approach incorporates both node and edge features, leveraging the complete spectrum of graph information. It employs the line graph transformation, where edges of the original graph are represented as nodes in a new graph, enabling effective learning of edge feature representations. Using two separate GCN encoders, the node features *X* and edge features *E* are encoded into latent spaces *Z*_*X*_ and *Z*_*E*_. Corresponding decoders then reconstruct the original features 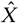 and Ê. The reconstruction process is optimized by minimizing a combined loss function based on the Frobenius norm, ensuring that the model captures the complex interactions inherent in the graph structure.

**Figure 2:**
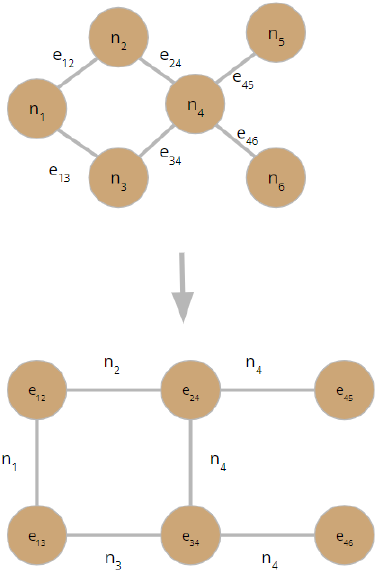
Dual of a Graph.

#### Algorithm 1

Dual Graph Autoencoding Framework

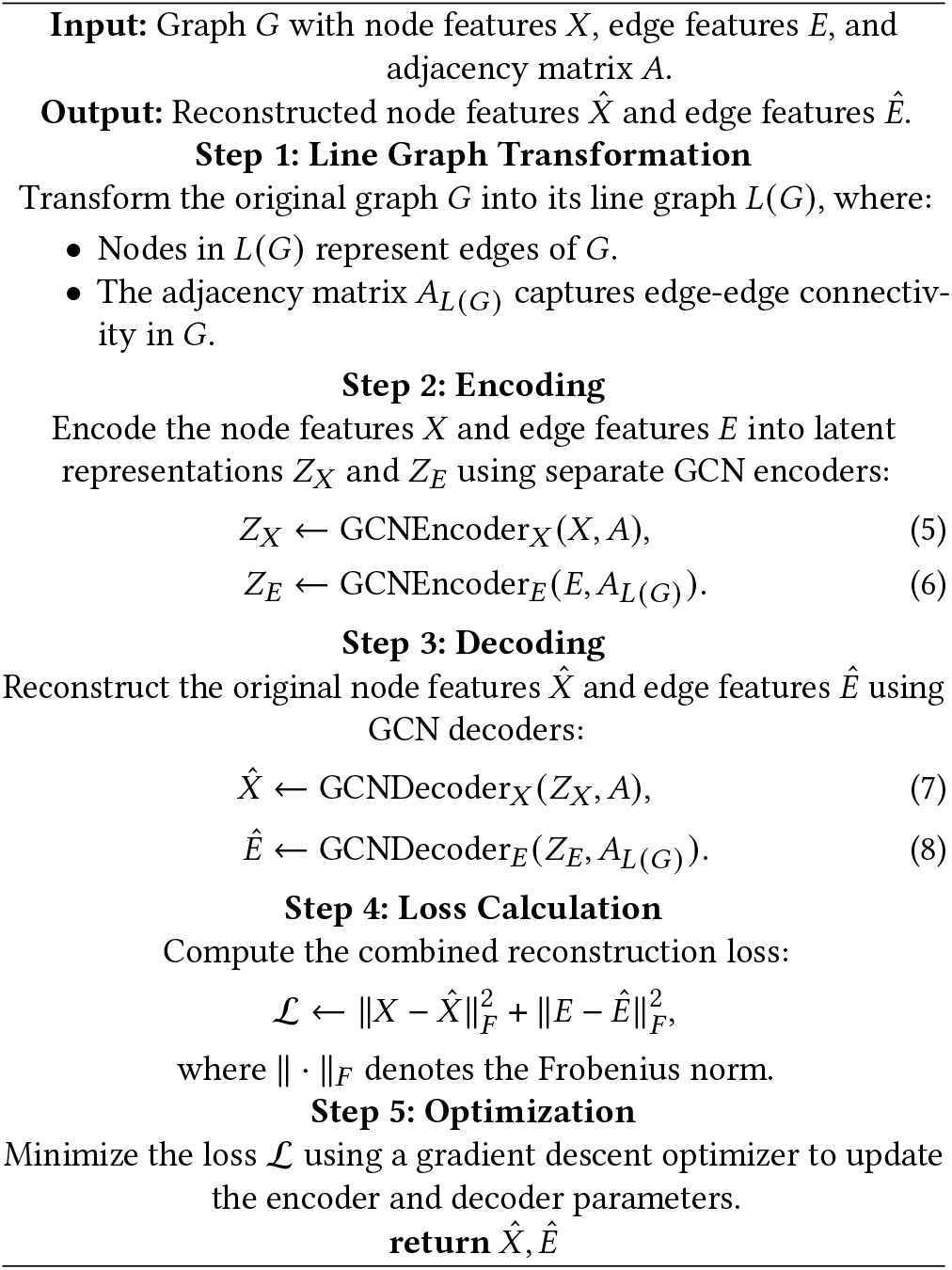

### 3.7 Complete Architecture and Downstream Network

The overall architecture of the proposed solution goes this way: we first use the constructed graph with rd-kit to encode it with the autoencoder and extract the hidden feature vector from the autoencoder. Then we convert the given graph into it’s dual graph, converting the edge features into node features and encode it using the autoencoder to provide another set of hidden features. Finally, we use chem-Bert to extract molecule level feature of size 768 and concatenate these three feature sets. In the downstream network use the the architecture shown in **??** to get a toxicity prediction for each datapoint from the consolidated feature sets as input. We used a combination of CNNs (Convolutional Neural Networks), Dense Layers and Dropout Layers. The Dense Layers are the backbone of any traditional NN. Including the CNN helps increase AUC-ROC score, since they can capture local clusters even in the embedding space. We train a separate model for each label, and use them for the corresponding prediction tasks. As such we end up with 12 separate models, with 2 autoencoders for each model (one for edge feature encoding and another for node feature encoding).

**Figure 3:**
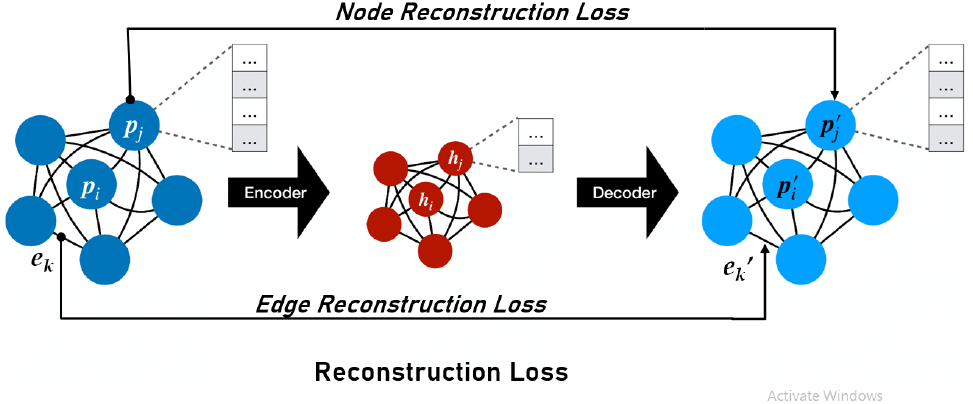
Loss functions.

## 4 RESULTS

### 4.1 Comparison with Different GNN Architectures

We first compare the performance of GraFusionNet with standard GNN based architectures. Particularly, we compare with four different approaches: node only, node-edge, language model features only, and graph transformer. Node only scheme uses the node features of the molecules derived from rdkit and employs a graph autoencoder to generate an embedding space of fixed size. Only edge scheme first converts the edge features of the graphs into node features with line graph conversion. This reconstructed graph is operated with the similar scheme to generate embedding vector. For language model based features we use the ChemBert features of the molecules and pass the embeddings through the prediction head. As GCN based graph neural network layers do not explicitly use edge features, we compare the performance of GraFusionNet with graphTransformers layer which also incorporates edge features during the GNN attention based concatenation step.

GraFusionNet demonstrates superior performance on the Tox21 dataset, achieving the highest average AUC-ROC score of 0.766. This improvement can be attributed to the effective incorporation of both node and edge features through the dual graph autoencoding framework. Notably, GraFusionNet achieves the best results in several classes, such as sr-mmp (0.888) and st-atad5 (0.799), indicating its ability to model complex relationships in molecular data. Compared to the Node Only Scheme, GraFusionNet improves the average score by 2.1%, underscoring the importance of leveraging edge features alongside node features. On the HIV dataset, GraFusionNet achieves a competitive score of 0.704. Although this is slightly lower than the Node Only Scheme’s score of 0.763, it highlights the framework’s adaptability across datasets. The slightly lower performance on the HIV dataset suggests that additional optimization may further enhance its generalization ability.

Table **??** highlights the overall effectiveness of GraFusionNet in handling molecular graph data. It consistently outperforms other methods on the Tox21 dataset and remains competitive on the HIV dataset. While language model features show strong results in certain cases, they fail to generalize as effectively. Graph Transformers, despite their theoretical advantages, perform poorly across both datasets, indicating a potential mismatch with the dataset properties or limitations in their design. These findings validate the robustness of the proposed dual graph autoencoding framework, especially in capturing both node and edge-level information for molecular property prediction.

### 4.2 Class-wise Performance Analysis

We evaluated the performance of various models on the Tox21 dataset, consisting of 12 binary classification tasks (e.g., nr-ahr, nr-ar, sr-p53) and the HIV task. This analysis focuses on the characteristics of each label and how different models perform on them.

The results reveal that certain tasks are consistently challenging across all models, while others show significant variation depending on the approach. For instance, the nr-ahr task sees the best performance from semantic models, with GraFusionNet achieving the highest score (0.876), closely followed by the Language Model Embedding (BERT) (0.873). This suggests that nr-ahr benefits heavily from semantic features, potentially due to underlying contextual patterns that are well-captured by BERT embeddings.

**Figure 4:**
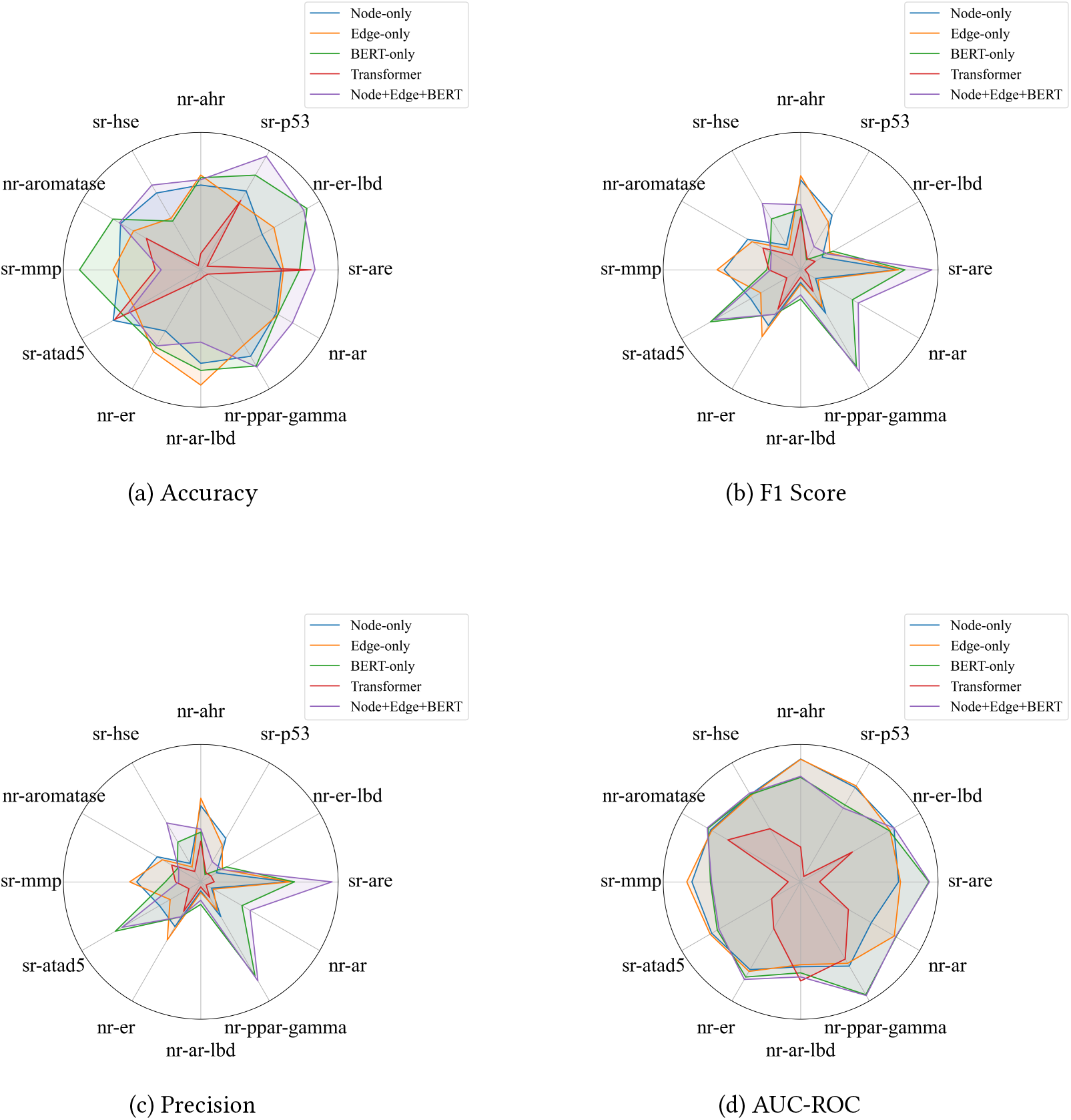
Metrics for the comparison of the different approaches studied.

In contrast, nr-ar-lbd, nr-aromatase, nr-ppar-gamma and sr-are stand out as tasks where structural features dominate. The GCN (Node Embedding) achieves the highest score in nr-aromatase (0.848), while the GCN (Node + Edge Embedding) excels in nr-ppargamma (0.805), nr-ar-lbd (0.790) and sr-are (0.760). These tasks likely rely on graph topology, where node connectivity and edge relationships play a crucial role. The underperformance of the BERT-only, GraFusionNet and transformer-based models on these labels further supports the importance of structural features.

The HIV task follows a similar trend. The GCN (Node + Edge Embedding) achieves the highest score (0.764), indicating the significance of structural features in capturing relationships critical to this dataset. While GraFusionNet achieves a competitive score (0.704), the added semantic information appears to introduce variability, making it less effective than the purely structural approach.

Some tasks, such as sr-mmp, demonstrate high variability in model performance. Here, GraFusionNet achieves the best score (0.888), followed closely by the Language Model Embedding (BERT) (0.883). This indicates that semantic information contributes significantly to this task, while structural features alone (as in GCN-based models) fail to achieve comparable results.

|  | Tox21 Classes |  |  |  |  |  |  |  |  |  |  |  |  | HIV Dataset |
| --- | --- | --- | --- | --- | --- | --- | --- | --- | --- | --- | --- | --- | --- | --- |
|  | nr-ahr | nr-ar | nr-ar-lbd | nr-atomatase | nr-er | nr-er-lbd | nr-ppar-gamma | sr-are | st-atad5 | sr-hse | sr-mmp | sr-p53 | Avg |  |
| **GCN (Node Embedding)** | 0.728 | 0.775 | 0.781 | **0.848** | 0.742 | 0.757 | 0.780 | 0.751 | 0.744 | 0.665 | 0.725 | 0.650 | 0.745 | 0.763 |
| **GCN (Node + Edge Embedding)** | 0.737 | 0.753 | **0.790** | 0.846 | 0.736 | 0.751 | **0.805** | **0.760** | 0.754 | 0.656 | 0.709 | 0.777 | 0.756 | 0.764 |
| **Language Model Embedding** | 0.873 | 0.750 | 0.684 | 0.759 | 0.742 | 0.772 | 0.691 | 0.720 | 0.785 | 0.694 | 0.883 | **0.787** | 0.761 | 0.634 |
| **Dual Graph Autoencoder + BERT** | **0.876** | **0.776** | 0.667 | 0.765 | **0.749** | **0.776** | 0.686 | 0.711 | **0.799** | 0.714 | **0.888** | 0.785 | 0.766 | 0.704 |
| **Graph Transformer with Edge** | 0.350 | 0.545 | 0.289 | 0.426 | 0.553 | 0.661 | 0.318 | 0.419 | 0.517 | **0.734** | 0.686 | 0.523 | 0.501 | 0.123 |

Interestingly, sr-hse highlights a unique case where the Graph Transformer with Edge achieves a relatively strong performance (0.734). This suggests that attention-based mechanisms may capture specific patterns in edge relationships that other models overlook. However, for most other tasks, this model struggles to generalize effectively.

Overall, the analysis reveals distinct patterns across tasks. Tasks like nr-ahr and sr-mmp favor semantic models, while tasks like nr-ar-lbd, nr-ppar-gamma, and HIV rely on structural features. Some tasks, such as sr-hse, show potential for hybrid approaches, leveraging attention mechanisms alongside structural information. These findings underscore the diversity of task requirements within the Tox21 and HIV datasets and the need for model designs tailored to specific label characteristics.

### 4.3 Interpretability Analysis

We visualized the t-SNE plots for each of our four approaches — **GraFusionNet** (node-edge-bert), node-edge, node only, and BERTonly — to explore their interpretability on the HIV dataset. These visualizations enabled us to analyze how each approach maps the data into a reduced-dimensional space, highlighting patterns and separability between positive and negative instances.

The node-only and node-edge approaches exhibit stronger clustering behavior compared to the other methods. In particular, we observe a prominent cluster of positive instances in the upper-right region of the t-SNE plots, corresponding to higher values along both dimensions. While these clusters indicate that the structural features of nodes and edges effectively capture relevant patterns in the dataset, some positive instances are still scattered across other regions. This suggests that while these approaches are robust in identifying positive samples, there is room for further refinement to achieve complete separation.

The BERT-only approach shows much weaker clustering. However, a small cluster of positive instances can be observed in the bottom-right region of the t-SNE plot. This pattern, though less distinct, suggests that BERT embeddings capture some meaningful semantic information about the dataset. Nonetheless, the lack of a more pronounced clustering indicates the limitations of using semantic features alone, particularly when structural information is crucial for effective modeling.

The **GraFusionNet** approach, which integrates both structural and semantic features, exhibits a small positive cluster to the right, beyond a certain radius in the t-SNE space. While this demonstrates the potential of combining node, edge, and BERT features, the clustering is weaker compared to the node-only and node-edge approaches. This may indicate that the integration of semantic features introduces additional variability, which can dilute the structural patterns that are more clearly observed in the simpler approaches.

In summary, the t-SNE analysis highlights the critical role of structural features—nodes and edges—in modeling the HIV dataset, as evidenced by the stronger clustering in the node-only and node-edge approaches. Although BERT embeddings add complementary semantic information, their standalone utility appears limited, and their integration in **GraFusionNet** requires further tuning to maximize their impact. These findings illustrate the nuanced trade-offs between structural and semantic features in graph-based learning tasks.

## 5 DISCUSSION AND FUTURE WORKS

In this research, we present not merely an alternative to the state-of-the-art (SOTA) approaches for graph neural networks (GNNs), but rather a shift in paradigm that emphasizes the significance of edge features within GNNs. Our proposed dual-graph autoencoder framework extends beyond the conventional node-centric GNN models by integrating edge features, thereby enriching the representational capacity of the network.

A noteworthy aspect of our study is the potential capability of our model to pass the Weisfeiler-Lehman (WL) graph isomorphism test, a benchmark that traditional GCNs struggle with due to their inherent limitations in encoding edge information. The incorporation of edge features might provide the means to discern non-isomorphic structures effectively, which standard GCNs typically fail to differentiate.

**Figure 5:**
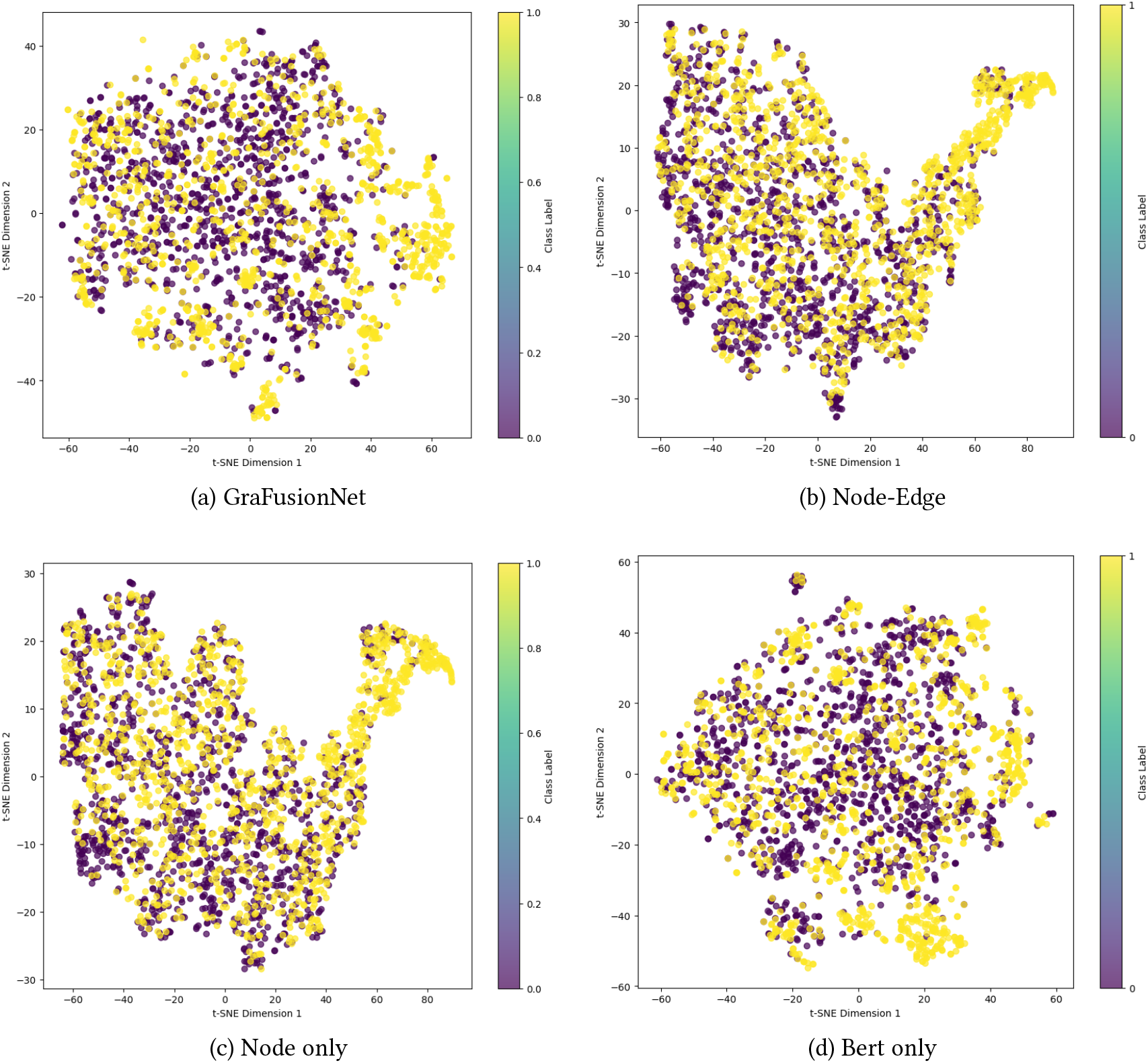
TSNE diagrams for each of our 4 approaches.

Looking ahead, several avenues open for exploration. Foremost among these is the need to validate the performance of our model across other standard datasets. Such an evaluation will not only fortify the generalizability of our approach but also establish a broader understanding of its effectiveness in varied contexts. Furthermore, a comparative analysis with transformer-based GNN models would offer insights into where our method stands concerning the latest advancements in GNN architectures (eg. GATConv, TransformerConv etc.)

In addition, no dataset-specific preprocessing was performed on our part. All other methods compared had ample steps in this regard. So incorporating a preprocessing step can improve the performance, we believe.

As we progress, it will also be crucial to examine the model’s scalability and how it performs with increasing graph sizes and complexity. Continuous refinement of the autoencoder’s architecture to efficiently process large-scale graphs while maintaining or enhancing its discriminative prowess will be key to its success.

In conclusion, our work lays down a foundational step towards a more holistic understanding of graph structures through GNNs, opening the door to deeper exploration and potential breakthroughs in graph analysis and interpretation.

## 6 CONCLUSION

In this paper, we have introduced a novel graph autoencoder network that leverages the dual graph concept to incorporate both node and edge feature reconstruction. Our approach represents a fundamental shift in graph neural network methodologies, where edge features have often been underutilized. By treating edges with equal importance as nodes, we have developed a more comprehensive model that captures the intricate relationships within graph data.

Our results indicate that this methodology not only provides a new lens through which to view graph-structured data but also outperforms many existing Graph Gated Neural Network (GGNN) based approaches. The effectiveness of our model is reflected in its superior performance on a variety of benchmark datasets, showcasing its potential to serve as a robust tool for graph analysis tasks.

As we conclude, it is worth emphasizing the significance of our approach as a step forward in graph representation learning. By integrating edge features into the autoencoding process, our model presents a more complete picture of graph data, leading to enhanced performance and opening new avenues for research and application. Moving forward, we anticipate that our contributions will inspire further studies, particularly in exploring the potential of edge feature reconstruction in graph neural networks.

